# *C. elegans* Nuclear Hormone Receptor NHR-49 promotes attractive chemotaxis independently of its role in fatty acid metabolism

**DOI:** 10.64898/2026.08.19.745339

**Authors:** James T.W. Wu, Glafira Ermakova, Junran Yan, Lexis D. Kepler, Mingzhe Estelle Zhang, Yiqing K. O. Wang, Yucheng Liu, Catharine H. Rankin, Stefan Taubert

**Author notes:** Current affiliation: Department of Biochemistry, University of Oxford, South Parks Road, Oxford OX1 3QU, The United Kingdom (Junran Yan).

## Abstract

All organisms must sense and adapt to environmental cues to survive and thrive. In the nematode worm *Caenorhabditis elegans*, the nuclear hormone receptor *nhr-49* is required for lipid homeostasis, stress resilience, pathogen defense, and longevity. In addition, recent studies suggest that NHR-49 controls avoidance behaviors and aversive memory to pathogenic bacteria and mitochondrial disruption, acting via its established role in fatty acid desaturation and fatty acid oxidation, respectively. However, the role of NHR-49 in neuronal signaling and behavior remains poorly understood. Here, we uncover a new role for NHR-49 in AWA neuron-mediated attractive chemotaxis. We show that *nhr-49* is required for chemotaxis towards diacetyl and pyrazine, two AWA neuron-sensed compounds, but not for AWC-or ASH-mediated chemotaxis. Supplementation with unsaturated fatty acids and mutational analysis of additional genes involved in fatty acid desaturation orβ-oxidation reveal that these processes do not affect attractive chemotaxis; this suggests that NHR-49 mediates attractive chemotaxis independently of its role in fatty acid desaturation and oxidation. Tissue-specific expression of NHR-49 in neurons or body wall muscle is sufficient to rescue chemotaxis defects, while *nhr-49* depletion in neurons alone causes chemotaxis defects. Finally, *nhr-49* mutants also show defects in another behavior, the response and habituation to mechanosensory stimuli. Together, our data reveal new functions for NHR-49 in sensory responses and non-associative learning, which are distinct from its roles in aversive behaviors and memory and, notably, independent of its well-established role in lipid metabolism.

**Significance statement:** Chemotaxis in the nematode worm *C. elegans* is critical for food finding and avoidance of harmful substances. Previous studies showed that nuclear receptor NHR-49 is involved in several avoidance paradigms in this model organism. These actions rely on lipid metabolism, a process directly regulated by NHR-49. The new study shows that NHR-49 is also critical for attraction to foo cues as mimicked by the chemical diacetyl. Unlike previous paradigms, this action is independent of lipid metabolism and appears to engage neuronal NHR-49 activity. Finally, *nhr-49* knockout mutants are also defective in habituation and memory formation. Collectively, this study describes new functions for NHR-49 in *C. elegans* and implies hitherto undiscovered roles in neuronal sensing and/or signalling.

## Introduction

All organisms must sense, respond to, and adapt to environmental cues to survive. Mounting responses to changing environments requires integrating sensory input with behavioral responses, which in turn requires coordinating nervous system activity with the function of other tissues. Dysregulation of the nervous system can impact the survival of animals living in the wild and lead to disease in humans. The nematode roundworm *Caenorhabditis elegans* is an ideal model for studying the mechanisms that underpin behaviors due to its small nervous system (∼300 neurons), short life cycle, and ease of genetic manipulation (Brenner, 1974). Despite its small nervous system, *C. elegans* demonstrates many fundamental behaviors that promote survival in response to a changing environment. In particular, *C. elegans* display complex chemotaxis behavior, with the ability to sense and move toward beneficial chemical stimuli like nutritional cues and away from potentially harmful stimuli like toxins or pathogenic microorganisms (Bargmann et al., 1993; Hilliard et al., 2002; Ferkey et al., 2021). *C. elegans* are also able to respond to mechanical stimuli. For instance, in response to a brief mechanosensory cue, *C. elegans* respond by reversing movement direction (Rankin et al., 1990; Goodman and Sengupta, 2019). Furthermore, *C. elegans* are capable of short-and long-term learning, including both associative and non-associative memory (Brandel-Ankrapp and Arey, 2023; Zhang et al., 2024). These animals also demonstrate habituation, a form of non-associative learning where there is decrease in response to a stimulus following multiple successive stimulations, to both olfactory and mechanical stimuli (Chiba and Rankin, 1990; Nuttley et al., 2001). Accordingly, work on *C. elegans* has revealed many new insights into the genetic regulation and molecular mechanisms that underpin olfaction, learning, and memory (Bargmann et al., 1993; Rose and Rankin, 2001).

The control of the aforementioned neuronal behaviours engages complex regulatory circuits, both cell autonomous and non-autonomous in nature. Recent evidence indicates that transcription factors of the nuclear hormone receptor gene family, which is greatly expanded in *C. elegans* versus mammals (Taubert et al., 2011), play important roles in this context. For example, NHR-76 reduces the expression of odorant receptors as worms age, thus promoting age-associated chemotaxis collapse (Yokosawa and Noma, 2025), whereas NHR-46 acts within neurons to control egg-laying behavior (Pender and Horvitz, 2018). Additionally, recent reports implicate NHR-49, best known for its control of lipid metabolism, longevity, and stress responses, in *C. elegans* neuronal and behavioural responses. Specifically, *nhr-49* is required for avoidance behaviors to pathogenic bacteria and for aversive memory in response to mitochondrial disruption (Kwon et al., 2023, 2024; Tsai et al., 2024). Notably, *nhr-49* regulates both of these activities through its control of various aspects of lipid homeostasis, namely fatty desaturation, fatty acid β-oxidation, and sphingolipid metabolism (Kwon et al., 2024; Tsai et al., 2024). This is reminiscent of NHR-49’s effect on other processes and phenotypes, such as the regulation of life span, fecundity, and innate immune responses (Gilst et al., 2005a, 2005b; Ratnappan et al., 2014; Naim et al., 2021), which also rely on the critical role of NHR-49 in lipid metabolism. Indeed, most NHR-49’s activities arise from its substantial impact on governing lipid metabolism, especially fatty acid desaturation.

However, recent studies provide emerging evidence that NHR-49’s functions extend beyond the regulation of lipid metabolism; for example, its role in the defence against pathogenic bacteria such as *Staphylococcus aureus, Pseudomonas aeruginosa,* and *Enterococcus faecalis* depends on both lipid metabolism genes as well as others, such as the flavin-containing monooxygenase 2 (*fmo-2*) (Dasgupta et al., 2020; Naim et al., 2021; Wani et al., 2021). Similarly, *nhr-49*-dependent regulation of lipid metabolism appears to play a minor role in hypoxia, where NHR-49 instead regulates autophagy genes to promote resilience (Doering et al., 2022). However, whether NHR-49 can influence neuronal or behavioural phenotypes independently of its impact on lipid metabolism is not known.

In line with the diversity of genes regulated by NHR-49, it functions in multiple tissues to control various phenotypes. For example, neuronal expression of NHR-49 restores *P. aeruginosa* survival and avoidance defects in the *nhr-49* null mutant (Naim et al., 2021; Kwon et al., 2023). In contrast, the short life span and the defects in hypoxia resilience of this mutant can be restored by *nhr-49* expression in multiple somatic tissues (Naim et al., 2021; Doering et al., 2022); however, the precise tissue(s) in which *nhr-49* is required for aversive memory in response to mitochondrial disruption is unknown. Therefore, although NHR-49 has established roles in metabolic and stress responses, whether it directly contributes to sensory processing and neuronal behavioral outputs remains poorly understood

Here, we investigated whether *nhr-49* plays a role in classical attraction chemotaxis, an essential behaviour when foraging for food. We show that *nhr-49* is essential for chemotaxis to compounds sensed by AWA neurons, but not those by AWC or ASH neurons. Oleic acid supplementation experiments and analysis of fatty acid metabolism defective mutants suggest that NHR-49 regulates chemotaxis independently of its role in fatty acid desaturation, distinguishing this role from previously observed *nhr-49*-dependent behaviors and memory. Finally, we show that NHR-49 regulates the response to mechanosensory stimuli and habituation to non-localized mechanosensory stimuli. Together, our data identify novel roles of NHR-49 in olfaction and sensory neuron signaling that are independent of its canonical role in lipid metabolism.

## Materials and Methods

### Nematode strains and growth conditions

We cultured *C. elegans* strains on nematode growth media (NGM) plates using standard techniques (Brenner, 1974). To minimize background genetic variation, each mutant was backcrossed into our lab wild-type N2 background at least six times. *E. coli* strain OP50 (CGC) was the food source in all experiments. All experiments were carried out at 20°C unless otherwise indicated. Worm strains used in this study are listed in Table 1. For synchronized worm growth, embryos were isolated by standard sodium hypochlorite treatment. Isolated embryos were allowed to hatch overnight on unseeded NGM plates until the population reached a synchronized halted development at the L1 stage via short-term fasting (12-24 hrs). Synchronized L1 stage larvae were then transferred to OP50-seeded plates and grown to the desired stage.

**Table 1.**
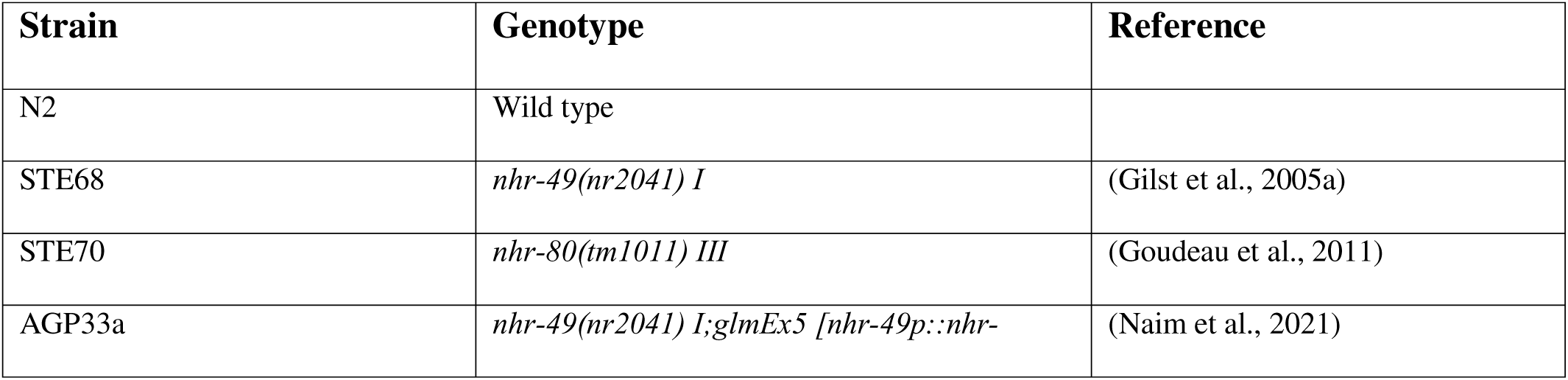
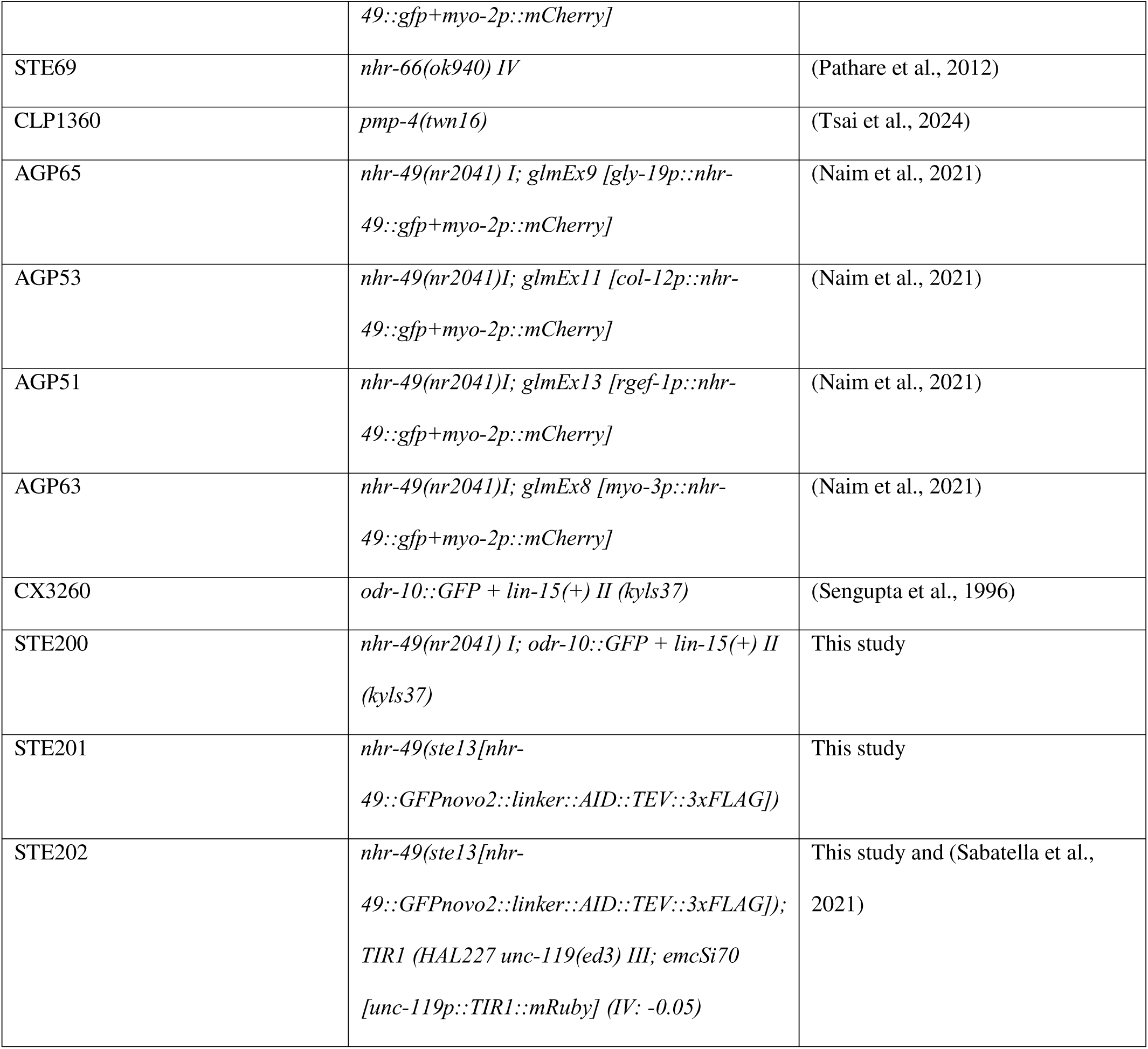
Worm strains used in this study.

We supplemented media with fatty acids as described (Deline et al., 2013) with minor changes. Briefly, NP-40 was added to liquid NGM agar at 65°C to a final concentration of 0.1%. Oleic acid (Sigma-Aldrich O1008) was then added to the media to a final concentration of 300µM or 3mM. Media plates were allowed to cool and then stored at room temperature in the dark to reduce fatty acid degradation for at least 2 and no more than 4 days before seeding with a 300µL overnight culture of OP50 grown in LB at 37°C. Seeded plates remained at room temperature and were used for experiments within 2 days.

### Generation of mutants with CRISPR-Cas9 genome editing

We used CRISPR-Cas9 genome editing to generate strain STE201(*nhr-49::GFPnovo2::AID*). We directly injected respective gRNA/Cas9 complexes (Cas9: IDT #1081059; tracrRNA: IDT #1072534) with the homology-directed repair (HDR) donor oligos, as described (Kurashina and Mizumoto, 2023). *myo-2p::mCherry* was used as co-injection marker. For endogenously tagging a previously published *nhr-49::GFPnovo2* strain (Tatge and Douglas, 2025) with the AID degron, we used two gRNAs surrounding the STOP codon of the endogenous *nhr-49::GFPnovo2* fusion. The AID sequence (from pJW1354: GSGGGG spacer-degron-TEV protease site-3x FLAG (Zhang et al., 2015)) was synthesized by Twist Biosciences (South San Francisco, CA, USA) and used directly as HDR repair template at a concentration of 10 ng/ul in the final injection mix. After injection, F1s bearing the co-injection marker were screened for heterozygous edits using PCR. Homozygous offspring of each line were confirmed by Sanger sequencing. gRNAs, HDR donor oligos, and genotyping primers are listed in the Table 2.

**Table 2.**
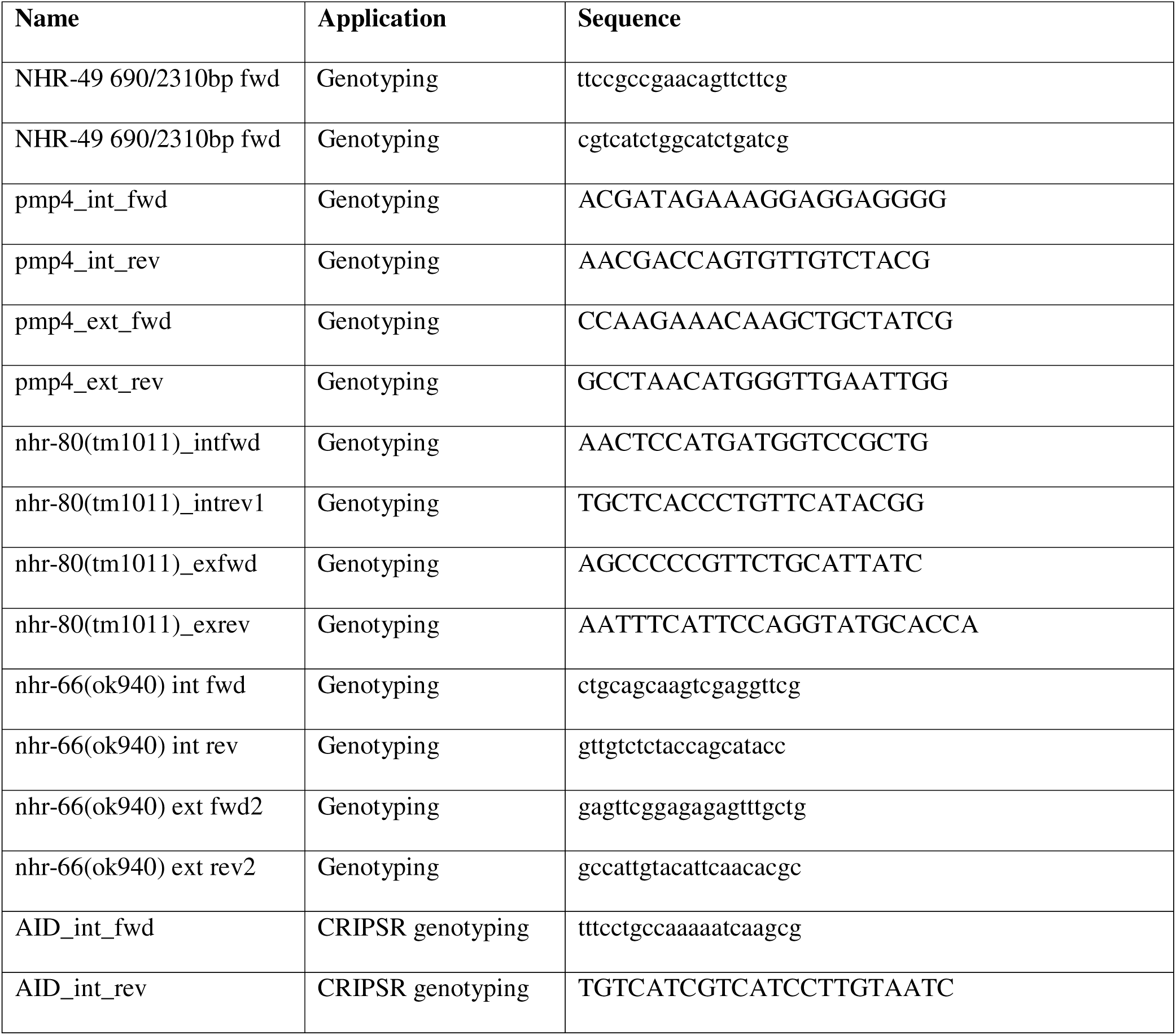
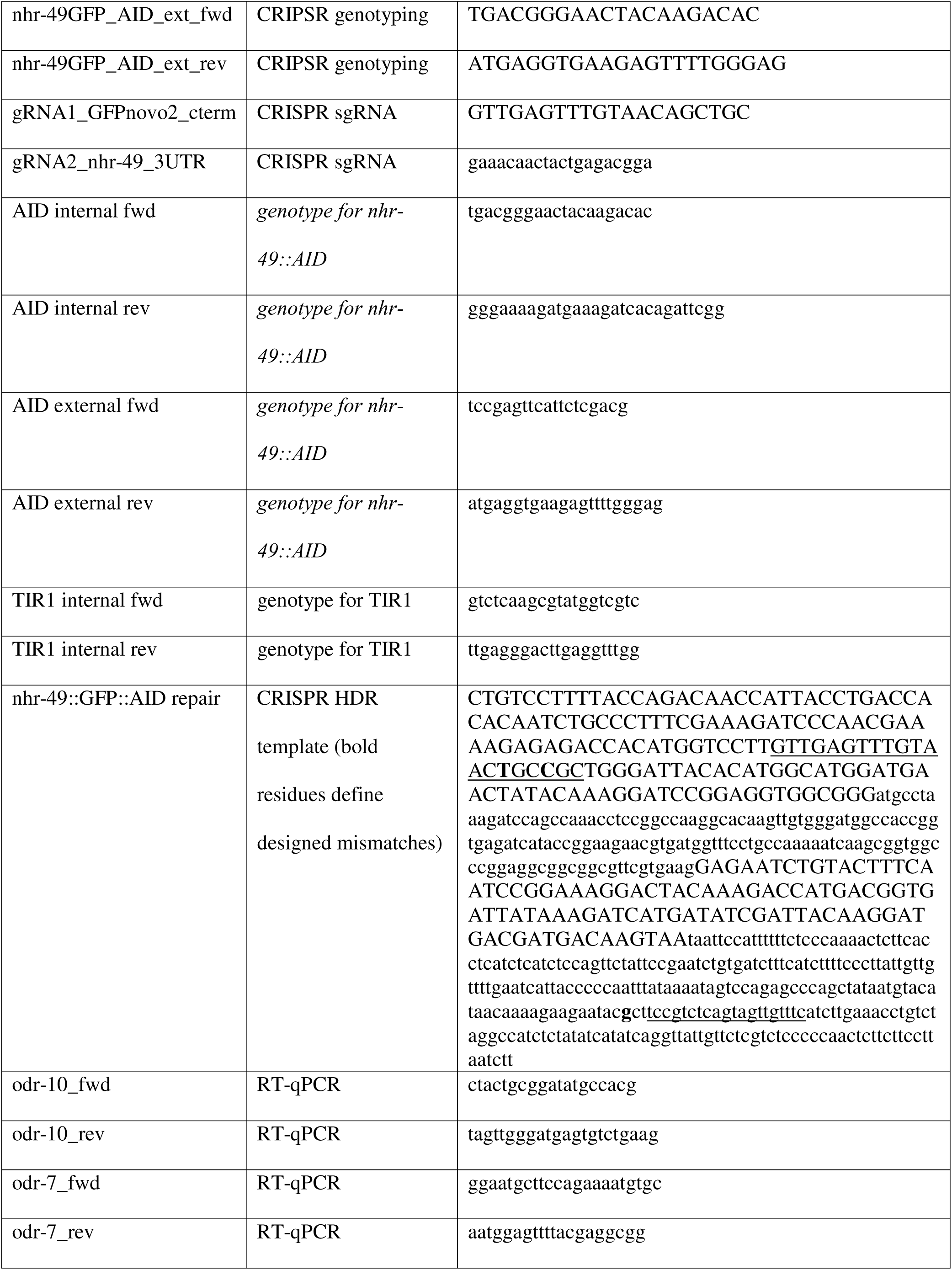
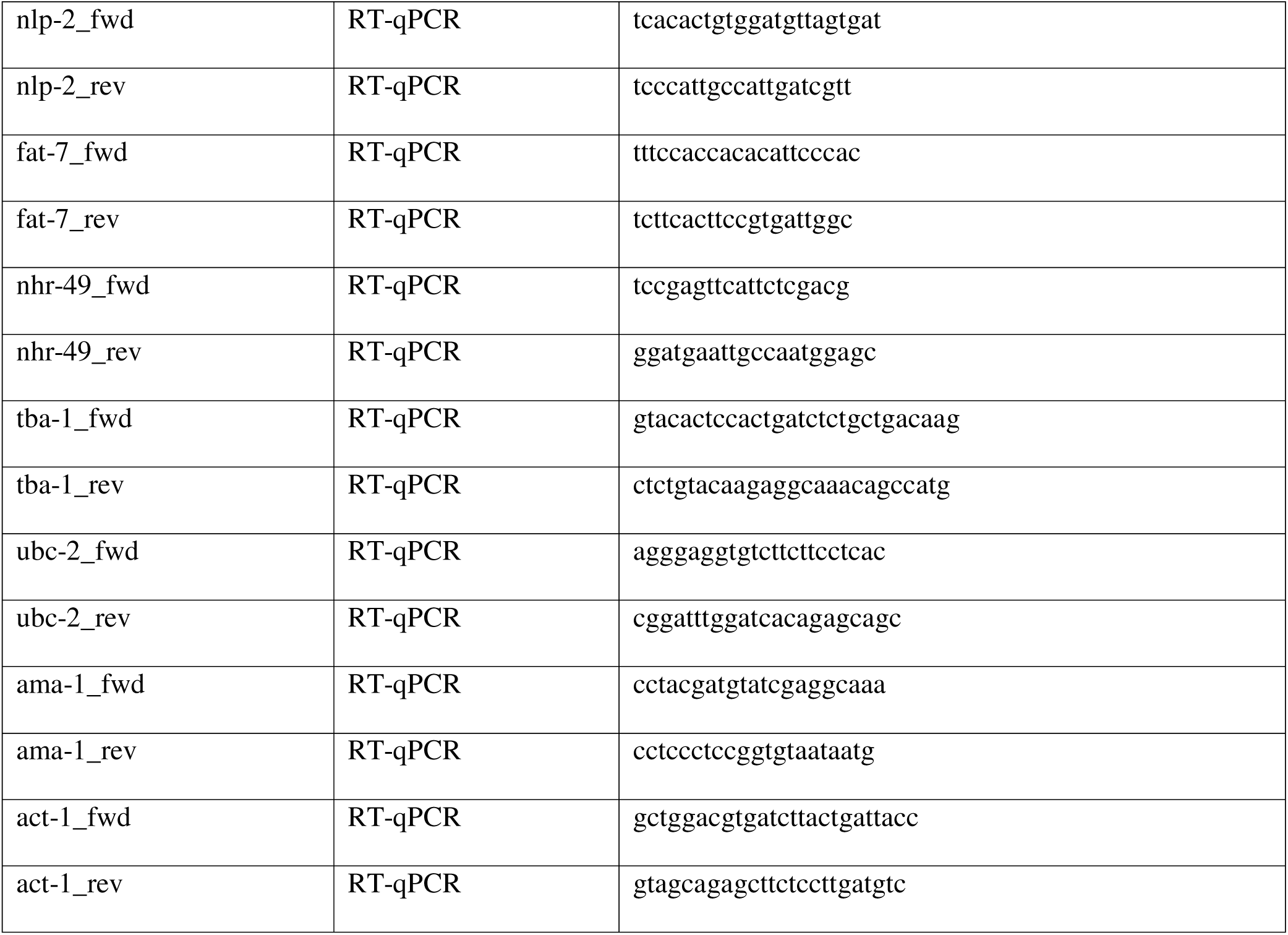
Primers used in this study.

### Chemotaxis assays

Chemotaxis assays were performed as described (Margie et al., 2013). Day 1 adult worms were collected with M9 buffer into 1.5mL centrifuge tubes. Pellets were washed with M9 at least three times to remove excess OP50. Each biological replicate contained three 6 cm NGM plates, each divided into 4 quadrants with a 1cm diameter circle in the middle, were used for each condition tested. Opposing quadrants on each plate were marked as test or control. 1µL of test solution was placed in the center of each test quadrant, and 1µL of 0.5M NaN_3_ in 50% ethanol was placed at the same spot. Test solutions were prepared as follows: 0.25% v/v diacetyl (Millipore Sigma #11038), 0.25% v/v isoamyl alcohol (Millipore Sigma #309435), 5mg/mL pyrazine (Millipore Sigma #P56003), or 50% v/v benzaldehyde (Millipore Sigma #B1334) with 0.25M NaN_3_ (Acros #19038-1000) in ethanol. Approximately 50-200 washed worms were pipetted into the center circle of each plate. Worms were left to move freely for 1 hour at 23°C and then counted. The chemotaxis index (CI) was calculated as follows: CI = (worms in test quadrants - worms in control quadrants) / (worms in test quadrants + worms in control quadrants). All worms in the 1cm diameter circle in the center of the plate were excluded from all calculations. Statistical comparisons between conditions were tested via Welch’s t-test.

For experiments with auxin-inducible degradation, chemotaxis assays were performed as described above with the following modifications. Age-synchronized populations of *nhr-49::AID*; *neuronal TIR1* (STE202) animals were generated by bleaching gravid adults. Synchronized L1 larvae were cultured at 20°C on NGML plates supplemented with either 100% ethanol as a vehicle control or 1 mM auxin (indole-3-acetic acid, Millipore Sigma I3750) dissolved in 100% ethanol to induce degradation of AID-tagged NHR-49. Day 1 adults were assayed. To maintain NHR-49 depletion throughout the chemotaxis assay, chemotaxis plates were prepared with the same ethanol or 1 mM auxin supplementation as the growth plates. Test/control quadrant preparation, data collection, and analysis were as described above.

### RNA isolation and RT-qPCR analysis

RNA was isolated from L4 stage worms as described (Yan et al., 2025). cDNA was generated from isolated RNA using Superscript III reverse transcriptase (ThermoFisher Scientific 18080093), random primers (ThermoFisher Scientific 48190011), dNTPs (Fermentas R0186), and RNaseOUT (ThermoFisher Scientific 10777019). Real-time quantitative PCR was conducted in 10µL reactions using Fast SYBR Master Mix (Life Technologies 4385612), undiluted cDNA, and 5µL primers. We analyzed the data with the ΔΔCt method. For each sample, we calculated normalization factors by averaging the (sample expression)/(average wild type expression) ratios of three normalization genes, *ama-1*, *tba-1*, and *ubc-2*. Average relative transcript abundance of normalization genes was kept at a constant across repeats to correct for batch variations in transcript abundance. The statistical significance of mRNA transcript level changes was calculated using a Welch’s t-test. Primers were tested with serial cDNA dilutions for PCR efficiency and are listed in Table 2.

### Analysis of fluorescent reporter lines via DIC and fluorescence microscopy

Day 1 adult worms were immobilized on 2% (w/v) agarose gel pads for microscopy using 0.05% levamisole (Sigma L9756). Images were captured at 63× magnification on Zeiss Axiocam 820 mono camera attached to a Zeiss Axioplan 2 compound microscope and acquired using ZEN 3.11 software (Zeiss). Images were collected using DIC and GFP fluorescence channels with identical exposure settings for all experiments (DIC: 198.362 ms, 70% illumination; EGFP: 500 ms, 70% illumination). Images were exported from ZEN 3.11 with scale bars for figure preparation.

### Behavioural profiling with Multi-Worm Tracker (MWT) assays

MWT experiments and analyses were performed as previously described (Kepler et al., 2022) with the following modifications. Age-synchronized populations were generated using a timed egg-lay protocol. Synchronization was performed on a dedicated set of MWT NGM agar plates prepared specifically for this purpose. These plates were seeded with 50 µL of *Escherichia coli* OP50 liquid culture, which was evenly distributed across the agar surface using sterile technique and a glass spreader to ensure uniform bacterial coverage. Plates were allowed to dry for at least 48 hours following pouring and an additional 48 hours after bacterial seeding.

For each genotype, six replicate synchronization plates were prepared. Five gravid hermaphrodites at 96 hours post-hatch were placed onto each plate and allowed to lay eggs for 4 hours before removal, yielding approximately 50-100 progeny per plate. On each experimental day, N2 wild type was synchronized in parallel with mutant strains under identical conditions. For each experiment, N2 was assayed alongside two to four mutant genotypes to enable direct, within-day comparison across strains.

Behavioural assays were conducted using MWT systems. For each experiment, six plate replicates per strain were assayed, with plates evenly distributed across three MWT units to control for instrument-specific variability. The order in which strains were tested was counterbalanced across the experimental session to minimize potential age-related effects arising from the duration of data collection.

Immediately prior to recording, Parafilm was removed from each plate, which was then placed onto the MWT stage. The lid was gently lifted to deliver a brief air puff (approximately 2-5 seconds) before the assay began. Each plate was subjected to a standardized 20-minute short-term habituation protocol. Animals were first allowed to acclimate to the recording environment for 5 minutes, followed by a 5-minute baseline recording period during which locomotion and morphological features were quantified in the absence of stimulation.

Following baseline data acquisition, animals received 30 non-localized mechanosensory stimuli delivered to the side of the plate using an automated push-solenoid at 10-second inter-stimulus intervals. These stimuli reliably elicited reversal responses, in which animals briefly moved backward before resuming forward locomotion. Multiple metrics of mechanosensory responsiveness and habituation learning were extracted from these reversal events, which reflect genetically dissociable underlying mechanisms. After the final stimulus, animals were given a 5-minute rest period, followed by delivery of a single additional stimulus to assess short-term retention of habituation (spontaneous recovery). All MWT assays were performed in a temperature-and humidity-controlled room maintained at approximately 20 °C and 40% relative humidity.

Stimulus delivery and image acquisition were controlled using MWT software (version 1.2.0.2). Behavioural features were quantified using Choreography software (version 1.3.0_r103552). Analyses were restricted to animals that moved more than two body lengths and were tracked for at least 20 seconds using the filters –shadowless, –minimum-move-body 2, and –minimum-time 20. Reversal events were identified using the MeasureReversal plugin, which detects animals initiating a reversal within 1 second of stimulus onset.

Choreography output files were processed and organized using custom R scripts. For each plate, behavioural metrics were calculated by averaging measurements across all tracked animals on that plate (typically 50-100 animals). Each plate was treated as a single biological replicate. For each genotype, plate-level values were averaged within each experimental day to generate a single daily mean. Daily genotype means were then compared across days using Welch’s t-test.

## Results

### *nhr-49* is required for attraction to AWA neuron-sensed molecules diacetyl and pyrazine

To test if NHR-49 plays a role in attractive chemotaxis, we assessed if an *nhr-49* mutation would affect the attraction of *C. elegans* to a well-studied odorant, diacetyl. Performing diacetyl chemotaxis assays using wild-type and *nhr-49(nr2041)* null mutant animals, we observed that loss of *nhr-49* substantially reduced attraction towards diacetyl at two different concentrations (Fig. 1A, B). Diacetyl is detected by AWA olfactory neurons, which also detect the odorant pyrazine (Sengupta et al., 1994). Thus, we also assessed pyrazine and observed that *nhr-49(nr2041)* mutants are also defective for attraction to pyrazine (Fig. 1C).

**Figure 1.**
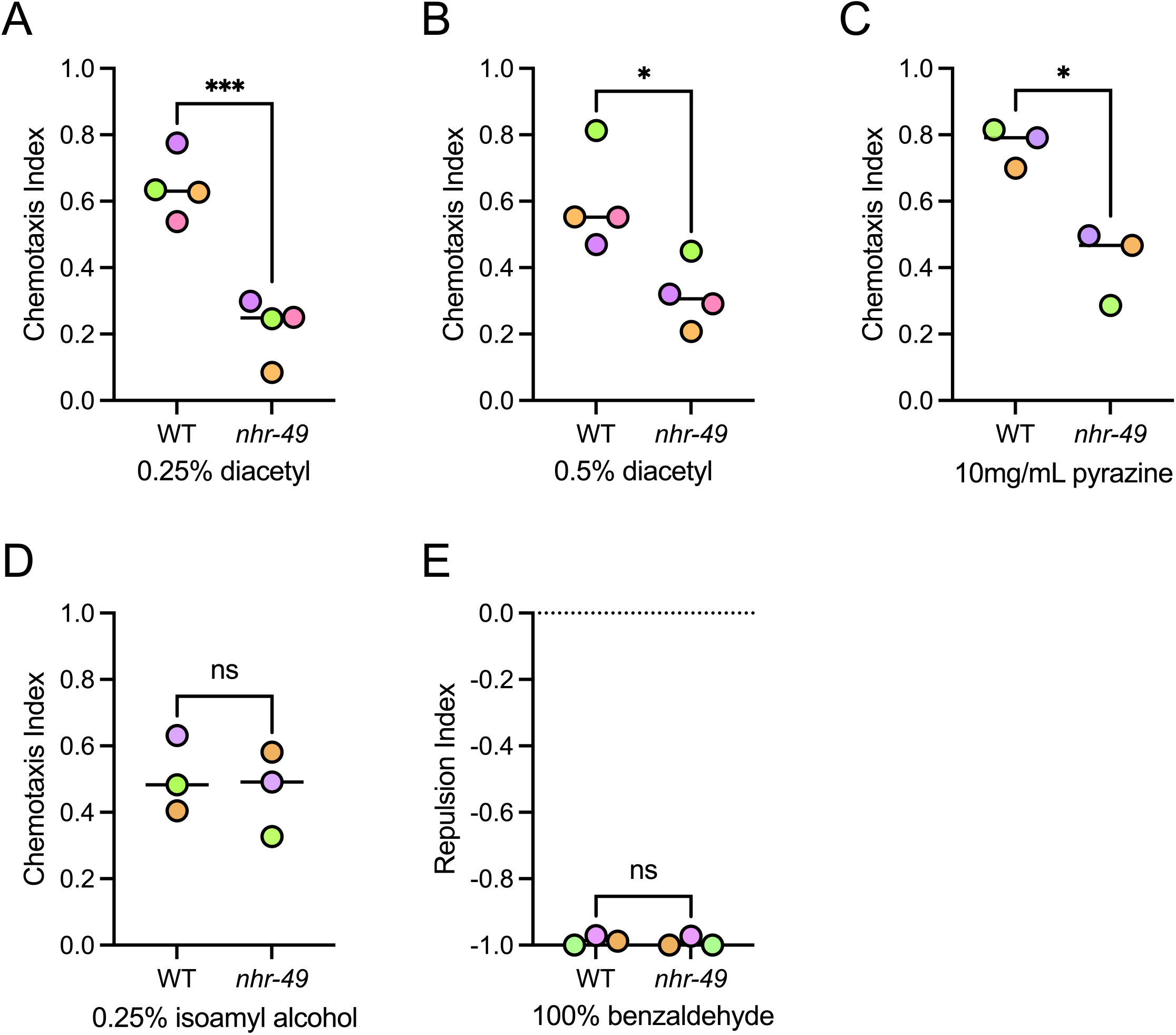
*nhr-49* is required for AWA neuron-mediated attractive chemotaxis. **(A)** Graph shows average chemotaxis index of wild-type (WT) and *nhr-49(nr2041)* null mutant worms towards 0.25% v/v diacetyl. N=4, p=0.0008, two-tailed Welch’s t-test. **(B)** Graph shows average chemotaxis index of WT and *nhr-49(nr2041)* null mutant worms towards 0.5% v/v diacetyl. N=4, p=0.025, two-tailed Welch’s t-test. **(C)** Graph shows average chemotaxis index of WT and *nhr-49(nr2041)* null mutant worms towards 10mg/mL pyrazine. N=3, p=0.0172, two-tailed Welch’s t-test. **(D)** Graph shows average chemotaxis index of WT and *nhr-49(nr2041)* null mutant worms towards 0.25% v/v isoamyl alcohol. N=3, p=0.7129, two-tailed Welch’s t-test. **(E)** Graph shows average repulsion index of WT and *nhr-49(nr2041)* null mutant worms to benzaldehyde. N=3, p=0.7252, two-tailed Welch’s t-test.

To test whether *nhr-49* loss generally affects chemoattractant sensing, we next assayed isoamyl alcohol, an attractive odorant that is specifically sensed by the AWC olfactory neurons (Colbert and Bargmann, 1995). In contrast to their defects in sensing diacetyl and pyrazine, *nhr-49(nr2041)* mutant animals exhibited no defects in attraction to isoamyl alcohol (Fig. 1D). Lastly, because *nhr-49* is implicated in aversive sensing (Kwon et al., 2024; Tsai et al., 2024), we tested whether the *nhr-49(nr2041)* is capable of sensing high concentrations benzaldehyde, which are detected in ASH neurons to trigger an avoidance behavior. As seen for isoamyl alcohol chemotaxis, we observed no defects of *nhr-49(nr2041)* mutant animals in benzaldehyde avoidance (Fig. 1E). We conclude that *nhr-49* plays a specific role in regulating attractive behaviors mediated by AWA neuron signaling, rather than being generally required for chemosensation.

### Lipid metabolic pathways regulated by NHR-49 are dispensable for AWA chemoattraction

*nhr-49*-dependent lipid metabolic processes are known drivers of neuronal sensing and signaling (Kwon et al., 2023; Tsai et al., 2024). For example, supplementation with oleic acid, an unsaturated fatty acid produced by the fatty acid desaturase *fat-7*, a strongly NHR-49-dependent gene, rescues the defects of *nhr-49* null mutant worms in pathogen avoidance (Kwon et al., 2024). To test if oleic acid is also involved in *nhr-49*-dependent diacetyl attraction, we supplemented wild-type and *nhr-49(nr2041)* mutant animals with oleic acid. We observed that this supplementation was unable to rescue the diacetyl chemotaxis defect of the *nhr-49(nr2041)* mutant (Fig. 2A). As another way to test if unsaturated fatty acids such as oleic acids are responsible for diacetyl sensing, we tested if worms mutant for *nhr-80*, which encodes a dimerization partner for NHR-49 that promotes the expression of *fat-7* and related genes *fat-6* and *fat-5* (Goudeau et al., 2011), also shows defects in diacetyl chemotaxis. Unlike *nhr-49(nr2041)* mutant animals, *nhr-80(tm1011)* mutant worms did not show defects in diacetyl attraction (Fig. 2B). Based on these data, we conclude that monounsaturated fatty acids are unlikely to be a key player in sensing or signaling attraction to diacetyl.

**Figure 2.**
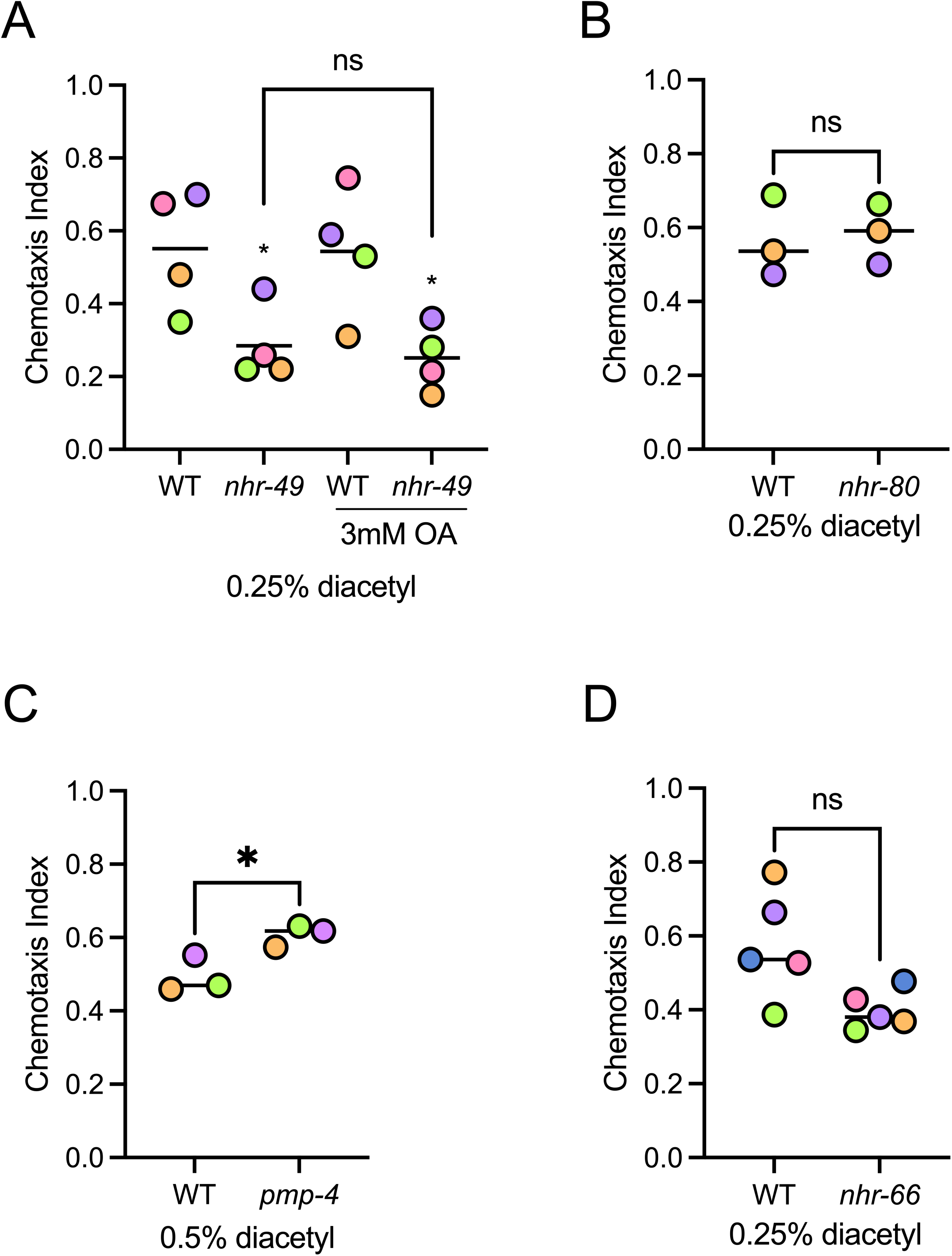
Role of lipid metabolism in AWA neuron-mediated attractive chemotaxis. **(A)** Graph shows average chemotaxis index of WT and *nhr-49(nr2041)* null mutant worms towards 0.25% v/v diacetyl on unsupplemented plates or on plates supplemented with 3mM oleic acid (OA). N=4, p=0.05 (WT vs. *nhr-49*) and 0.0242 (WT + OA vs. *nhr-49* + OA), ordinary one-way ANOVA with Dunnett’s multiple comparison test. **(B)** Graph shows average chemotaxis index of WT and *nhr-80(tm1011)* null mutant worms towards 0.25% v/v diacetyl. N=3, p=0.8222, two-tailed Welch’s t-test. **(C)** Graph shows average chemotaxis index of WT and *pmp-4(twn16)* null mutant worms towards 0.5% v/v diacetyl. N=3, p=0.0395, two-tailed Welch’s t-test. **(D)** Graph shows average chemotaxis index of WT and *nhr-66(ok940)* null mutant worms towards 0.25% v/v diacetyl. N=5, p=0.0509, two-tailed Welch’s t-test.

Another recent study showed that aversive memory triggered by mitochondrial disruption involves *nhr-49* and genes regulated by NHR-49 involved in peroxisomal β-oxidation, such as *pmp-4*, *dhs-23*, and *daf-22* (Tsai et al., 2024). To test if a similar mechanism is involved in diacetyl attraction, we assessed *pmp-4(twn16)* mutants. Strikingly, *pmp-4(twn16)* mutant animals were no worse than wild type in their attraction to diacetyl (Fig. 2C). Finally, NHR-66 dimerizes with NHR-49 to specifically express genes involved in sphingolipid metabolism (Pathare et al., 2012), but its role in neuronal sensing and signaling has not yet been studied. We assessed the ability of *nhr-66(ok940)* null mutants in the diacetyl attraction paradigm and found that these worms displayed a mildly reduced attraction to diacetyl when compared to wild type that approached significance (Fig. 2D). We conclude that fatty acid desaturation and peroxisomal β-oxidation are unlikely to be involved in diacetyl attraction, and that NHR-49 likely regulates this sensing and signaling process via a new mechanism, possibly involving sphingolipid metabolism.

### Neurons and muscle are involved in *nhr-49*-mediated chemotaxis

NHR-49 is expressed broadly, including in the intestine, neurons, muscle, and hypodermis (Gilst et al., 2005a; Naim et al., 2021), and can act in multiple tissues to promote resistance to stress and pathogen infection, whereas its pro-longevity role may more specifically depend on neurons (Naim et al., 2021; Doering et al., 2022). To test which tissues NHR-49 acts in to promote diacetyl chemotaxis, we studied strains that express an NHR-49::GFP translational fusion protein in the *nhr-49(nr2041)* mutant background from tissue-specific promoters (Naim et al., 2021). Comparing these rescue strains to their non-GFP siblings, respectively, we found that expressing NHR-49::GFP from its own promoter was sufficient to restore attraction to diacetyl (Fig. 3A). Moreover, expression of NHR-49::GFP from neuron-and body wall muscle-specific promoters also rescued chemotaxis, whereas expression from hypodermis-or intestine-specific promoters was not sufficient to promote chemotaxis (Fig. 3B-E). We conclude that NHR-49 activity in the neurons or muscle is sufficient to restore diacetyl chemotaxis.

**Figure 3.**
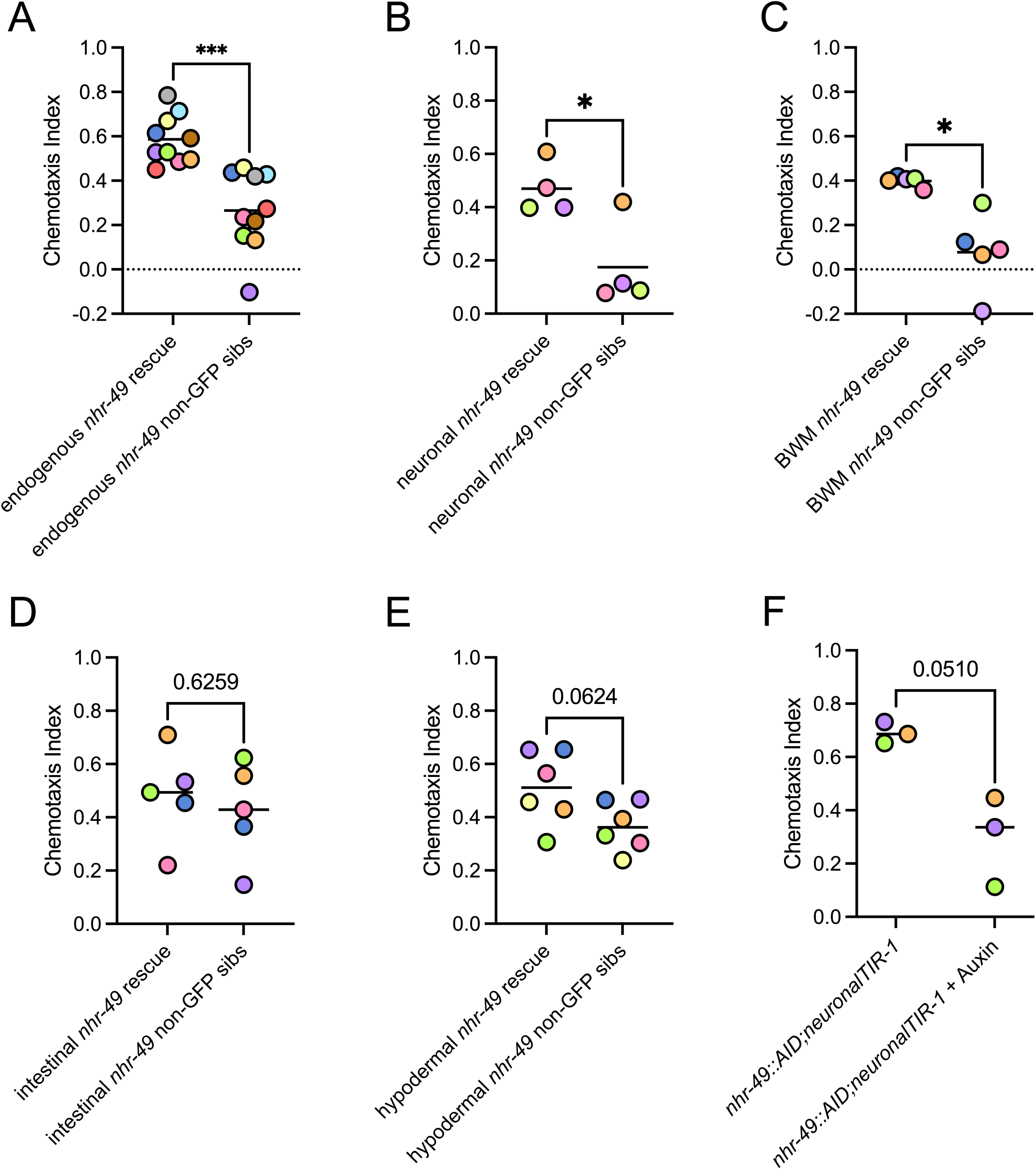
Analysis of tissue-specific function of NHR-49 in attractive chemotaxis. **(A)** Graph shows average chemotaxis index of *nhr-49(nr2041)* null mutant worms expressing NHR-49::GFP from the *nhr-49* promoter and non-transgenic siblings towards 0.25% v/v diacetyl. N=10, p=0.0002, two-tailed Welch’s t-test. **(B)** Graph shows average chemotaxis index of *nhr-49(nr2041)* null mutant worms expressing NHR-49::GFP from the neuronal *rgef-1* promoter and non-transgenic siblings towards 0.25% v/v diacetyl. N=4, p=0.0281, two-tailed Welch’s t-test. **(C)** Graph shows average chemotaxis index of *nhr-49(nr2041)* null mutant worms expressing NHR-49::GFP from the body-wall muscle promoter *myo-3* and non-transgenic siblings towards 0.25% v/v diacetyl. N=5, p=0.0141, two-tailed Welch’s t-test. **(D)** Graph shows average chemotaxis index of *nhr-49(nr2041)* null mutant worms expressing NHR-49::GFP from the intestinal promoter *gly-19* and non-transgenic siblings towards 0.25% v/v diacetyl. N=5, p=0.6259, two-tailed Welch’s t-test. **(E)** Graph shows average chemotaxis index of *nhr-49(nr2041)* null mutant worms expressing NHR-49::GFP from the hypodermal promoter *col-12* and non-transgenic siblings towards 0.25% v/v diacetyl. N=6, p=0.0624, two-tailed Welch’s t-test. **(F)** Graph shows average chemotaxis index of NHR-49::GFP::AID;*unc-119p*::TIR1 worms without and with 1mM auxin added towards 0.25% v/v diacetyl. N=3, p=0.051, two-tailed Welch’s t-test.

The tissues specific rescue experiments suggested that *nhr-49* function in neurons is sufficient to promote chemotaxis. To test if it is required in this tissue, we used the auxin inducible degradation (AID) system to selectively deplete *nhr-49* in neurons. We generated an NHR-49::AID strain and crossed into worms expressing a neuron-specific TIR1 degradation driver (Sabatella et al., 2021). In these worms, administration of auxin resulted in a substantial and borderline significant reduction of chemotaxis (Fig. 3F). This demonstrates that neuronal *nhr-49* plays important roles in attractive chemotaxis.

### *nhr-49* is required to express the AWA-specific *odr-10p::gfp* reporter

Attraction to diacetyl and pyrazine is controlled by the AWA neurons and involves the activity of the AWA-resident transcription factor ODR-7 and its target ODR-10, a G protein-coupled olfactory receptor (Sengupta et al., 1996). To test if *nhr-49* affects the expression of these genes and the Neuropeptide-Like Protein NLP-2, which regulates sleep and wakefulness and also shows AWA-enriched expression (Van der Auwera et al., 2020), we quantified the mRNA levels of these neuronal marker genes with RT-qPCR. In whole-animal RNA extracts, loss of *nhr-49* did not cause downregulation of *odr-7* or *odr-10* but did downregulate *nlp-2* expression, albeit modestly Fig. 4A; in comparison, the well-characterized NHR-49 regulated gene *fat-7* (Gilst et al., 2005a, 2005b) showed an almost complete loss of expression in *nhr-49(nr2041)* null mutant animals (Fig. 4A). *odr-7*, *odr-10*, and *nlp-2* are expressed in tissues and cell types other than the AWA neurons, such as the body wall muscle, coelomocyte, intestine, hypodermis, and germ line (Hammarlund et al., 2018), confounding insights on AWA-specific expression in whole animals by RT-qPCR. Because whole-animal RT-qPCR cannot distinguish expression within AWA neurons from expression in other tissues, we next examined *odr-10* expression using an AWA-specific GFP reporter. To assess if *nhr-49* loss affects *odr-10* expression in AWA neurons, we crossed the *nhr-49(nr2041)* null allele into a strain bearing an *odr-10p::gfp* transcriptional reporter; these worms exhibit strong and specific GFP expression in the AWA neuron (Sengupta et al., 1996). Notably, *nhr-49* mutation substantially reduced the proportion of worms with detectable GFP signal (Fig. 4B,C). These findings suggest that *nhr-49* is required for normal *odr-10p::gfp* expression in AWA neurons, which may reflect effects on neuronal development, cell identity, or *odr-10* transcriptional regulation.

**Figure 4.**
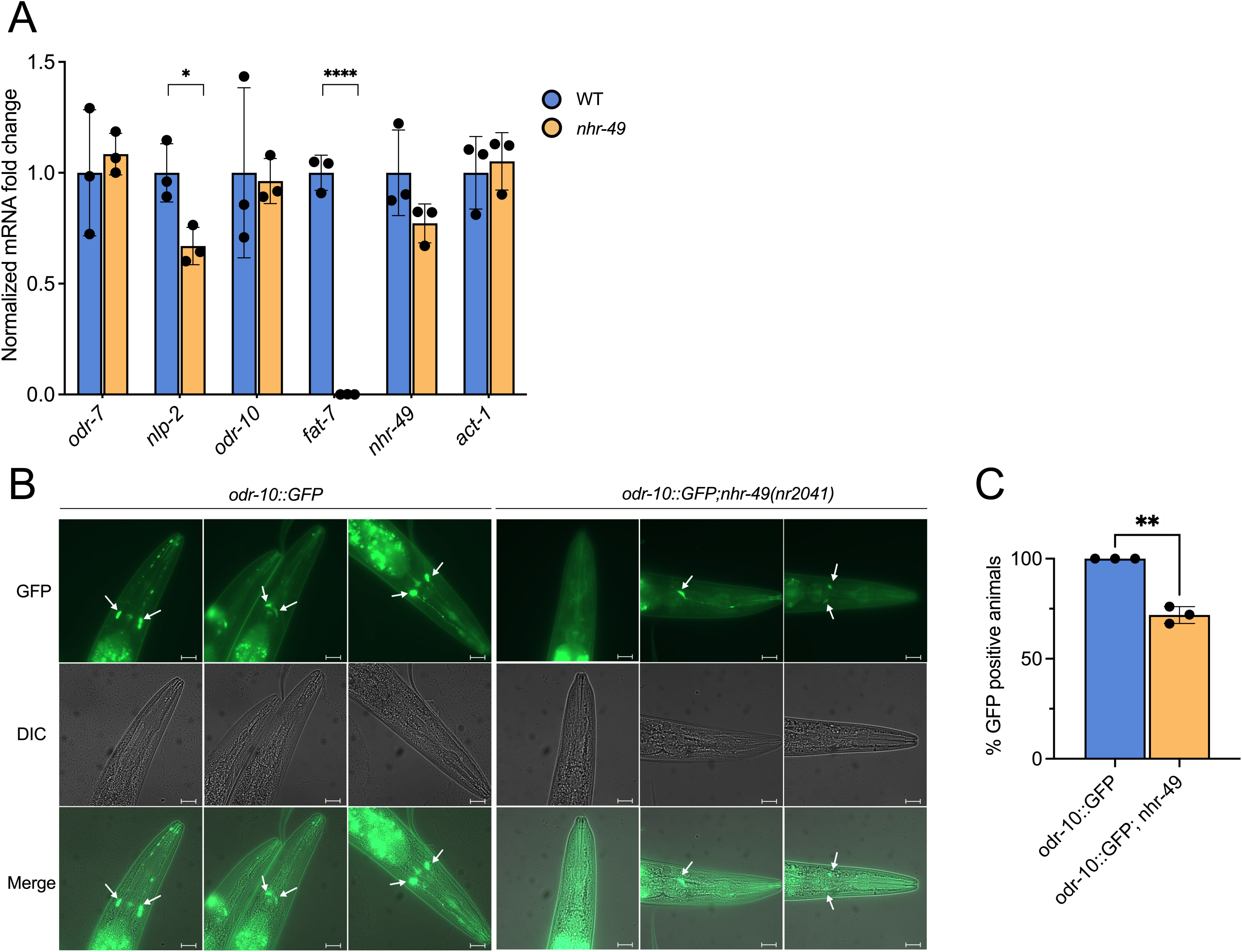
*nhr-49* null mutants show defects in markers specific for the AWA neuron. **(A)** Representative images of the GFP signal from an *odr-10p::gfp* transcriptional reporter. Size bar represents 20μm. White arrows point to GFP-positive AWA neuron-specific GFP signal. **(B)** Quantification of the GFP signal from an *odr-10p::gfp* transcriptional reporter. Quantification shows the fraction of animals that show at least one GFP-positive AWA neuron; N=3 experiments of 25-35 animals, **p= 0.0074, two-tailed Welch’s t-test. **(C)** Real-time quantitative PCR (RT-qPCR) of *odr-7*, *nlp-2*, *odr-10*, *fat-7*, *nhr-49*, and *act-1* mRNA expression in WT and *nhr-49(nr2041)* mutants, normalized to *ama-1* mRNA. N=4; p=0.6654 (*odr-7*), 0.0284 (*nlp-2*), 0.8847 (*odr-10*), 0.0021 (*fat-7*), 0.1661 (*nhr-49*), and 0.6904 (*act-1*), two-tailed Welch’s t-test.

### Loss of NHR-49 alters mechanosensory responses and habituation

Because our data show that NHR-49 plays a role in the sensing or signaling in the presence of volatile molecules, we hypothesized that NHR-49 might be more broadly involved in neuronal sensation/signalling processes. In line with broad roles for NHR-49 in neurons, its mRNA is detectable in many cells of the *C. elegans* nervous system, as evidenced by CenGEN data (Hammarlund et al., 2018).

To gain further insight into NHR-49’s role in neuronal function, we used a machine vision tracking system, the Multi-Worm Tracker (MWT) (Swierczek et al., 2011). Because *nhr-49* is expressed in neurons mediating mechanosensory behaviours (Hammarlund et al., 2018), we used the MWT to examine the effect of *nhr-49* loss on the tap withdrawal response, i.e., the temporary reversal of animals upon delivery of a non-localized mechanosensory stimulus (the tap) (Rankin et al., 1990). In addition to assessing mechanosensation, this allowed us to assess *nhr-49*’s role in habituation (determined by assessing animal response to 30 consecutive mechanosensory stimuli given at 10 second inter-stimulus intervals) and short-term memory (assessed by delivering a 31^st^ stimulus five minutes after the 30^th^ stimulus to assess retention of the habituated response level). As previously reported (Kwon et al., 2023), loss of *nhr-49* caused decreased animal size, specifically width, length, and overall area, as well as on-food baseline speed (Fig. 5A-D). *nhr-49* mutants were as likely as wild type to respond to mechanosensory stimulation (Fig. 5E), but displayed a shorter reversal distance and reduced speed after the first stimulus (Fig. 5F, G). This suggests that *nhr-49* does not impact the overall responsivity to a single stimulus while the observed defects likely reflect its role in maintaining overall movement velocity as seen at baseline (Fig. 5D). Interestingly, *nhr-49* deletion affected mechanosensory habituation, as the difference in response probability from the first to the final three stimuli was lower compared to wild type (Fig. 5H,I), whereas *nhr-49* loss did not affect short-term memory of reversal probability (Fig. 5J). Thus, *nhr-49* is dispensable for initial stimulus detection and memory formation but required for habituation to persistent mechanical stimulation. Together, these data validate known developmental and movement phenotypes of *nhr-49* mutants and suggest a novel role for NHR-49 in mechanosensation and habituation.

**Figure 5.**
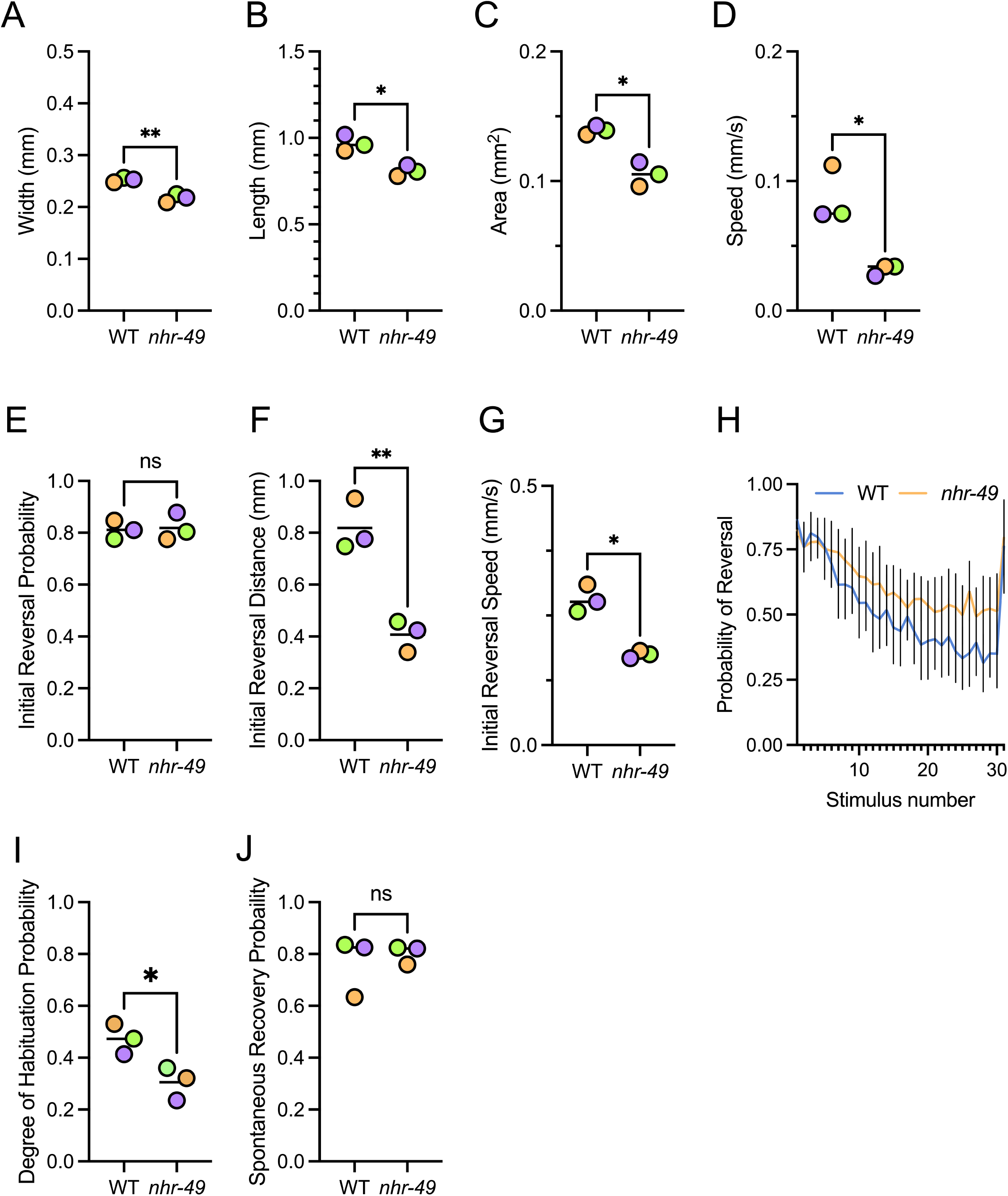
Mechanosensation and habituation defects in *nhr-49* null mutants. (A-D) Graphs show wild type and *nhr-49* mutants’ baseline **(A)** width, **(B)** length, **(C)** area, and **(D)** speed. **(E-G)** After initial mechanical stimulus: graphs show measurements of wild type and *nhr-49* mutants’ initial **(E)** reversal probability, **(F)** reversal distance, and **(G)** reversal speed. **(H-J)** After repeated mechanical stimuli: graphs show **(H)** overall habitation curve for reversal probability and **(I)** degree of habituation probability, and **(J)** spontaneous recovery of reversal probability. For all experiments n=3, two-tailed Welch’s t-test; p= 0.0060 **(A)**, 0.0114 **(B)**, 0.0155 **(C)**, 0.0121 **(D)**, 0.8434 **(E)**, 0.0064 **(F)**, 0.015 **(G)**, 0.0291 **(I)**, and 0.6366 **(J)**.

## Discussion

The *C. elegans* nuclear hormone receptor NHR-49, a sequence and functional ortholog of mammalian Peroxisome Proliferator-Activated Receptor alpha (PPARα) and Hepatocyte Nuclear Factor 4-alpha (HNF4α), plays important roles in the regulation of metabolism, stress resilience, response to pathogen infection, and life span (Doering et al., 2023). More recently, new roles for NHR-49 in the control of neuronal behaviors have emerged, including in egg laying, pathogen avoidance, and aversive memory establishment caused by mitochondrial disruption (Pender and Horvitz, 2018; Kwon et al., 2023, 2024; Tsai et al., 2024). These behaviors have been linked to NHR-49’s well-stablished regulation of fatty acid desaturation and β-oxidation (Kwon et al., 2024; Tsai et al., 2024), respectively, suggesting that these behavioral defects are downstream consequences of defective fatty acid metabolism, consistent with existing paradigms. In contrast, our present study reveals a new behavior under NHR-49’s control, AWA neuron-controlled attractive chemotaxis, that appears to be independent of its canonical role in fatty acid desaturation and β-oxidation and instead may involve other downstream mechanisms, potentially including sphingolipid metabolism. Our study therefore expands the repertoire of behaviors directed by NHR-49 and illuminates potential new mechanisms by which this transcription factor regulates neuronal function.

### A new role for NHR-49 in attractive chemotaxis

Recent studies have delineated several functions for NHR-49 that extend beyond its control of metabolism, stress resilience, and aging. Specifically, NHR-49 promotes pathogen avoidance to *P. aeruginosa* via its known regulation of fatty acid desaturation, and enhances short-term avoidance memory during oxidative stress by regulating peroxisomal β-oxidation (Kwon et al., 2024; Tsai et al., 2024). Our work substantially expands on these findings. First, we show that *nhr-49* is required for the chemotaxis to attractive compounds sensed by AWA, but not by AWC and ASH neurons. That NHR-49 is involved in detecting food cues is especially intriguing as NHR-49 promotes the expression of fatty acid β-oxidation genes, especially in starvation (Gilst et al., 2005b, 2005a). Integrating the detection of food cues with the control over access to stored nutrients into one regulator is evolutionarily plausible and could reflect a conserved strategy to achieve behavioral and metabolic coordination, thus optimizing chances for survival.

Second, the role of NHR-49 in attractive chemotaxis appears to be independent of its known effects on fatty acid desaturation and β-oxidation. Lipid molecules are key mediators of chemotaxis in *C. elegans* (O’Halloran et al., 2009). Polyunsaturated fatty acids (PUFAs) are involved in neuronal signaling, and mutants for the fatty acid desaturase genes *fat-1*, *fat-3*, and *fat-4* are defective in chemotaxis and nociception (Kahn-Kirby et al., 2004). TRPV channels, such as OSM-9 and OCR-2, are thought to require PUFAs for activation and chemotaxis responses (O’Halloran et al., 2009). Notably, the AWA neurons, which control chemotaxis to diacetyl and pyrazine, express both OSM-9 and OCR-2 (Hammarlund et al., 2018). NHR-49 is a key regulator of lipid metabolism, upregulating the fatty acid desaturase genes *fat-5* and *fat-7* (Gilst et al., 2005a, 2005b), which promote the production of monounsaturated fatty acids (Watts and Ristow, 2017). We therefore used complementary genetic and dietary approaches to test whether fatty acids might act downstream of NHR-49 in diacetyl attraction. First, we tested genetic mutants for both regulators (*nhr-80*, *nhr-66*) and effectors (*pmp-4*, the same mutant that showed defects in aversive memory formation) of fatty acid metabolism. Second, we directly tested whether fatty acid supplementation is sufficient to rescue defects in attractive chemotaxis. Interestingly, both approaches consistently showed that diacetyl chemotaxis is regulated independently of established, *nhr-49*-mediated fatty acid desaturation or oxidation pathways, suggesting that NHR-49 may engage different, novel mechanisms and processes to regulate this specific behavior. Our data suggest that NHR-66, which dimerizes with NHR-49 to control expression of sphingolipid metabolism genes, may contribute to AWA-driven chemosensation. However, further investigation into the role of *nhr-66* and/or sphingolipid metabolism will be required to conclusively determine their contribution.

### Tissue specificity of NHR-49 action

NHR-49 exerts its functional roles in multiple tissues, including the neurons, intestine, hypodermis, and others. Rescue experiments of *nhr-49* null mutants have revealed an essential role for neuronal *nhr-49* in other behaviours: for example, restoration of *nhr-49* expression in cholinergic and glutamatergic neurons was sufficient to rescue avoidance of pathogenic bacteria, whereas expression in cholinergic and serotonergic neurons effectively restored life span (Kwon et al., 2023, 2024). In our study, AWA-associated chemotaxis behavior of *nhr-49* mutants was restored by expression of NHR-49 from pan-neuronal and body-wall muscle promoters; the latter implies that the observed phenotypes may arise due to defects in signaling pathways engaging endocrine or metabolic molecules. We note that rescue of mutants with overexpression from strong, tissue-specific promoters may lead to artifacts and non-tissue-specific expression. However, an important role for NHR-49 within neurons is cemented by the fact that depletion specifically in this tissue results in substantial chemotaxis defects. Future studies defining the neuron-and AWA-specific transcriptomes under the influence of NHR-49 would be insightful.

### NHR-49 impacts the AWA neuron chemoreceptor ODR-10

The transcription factor ODR-7 and its target, the GPCR ODR-10, are critical players in establishing AWA neuron differentiation and function (Sengupta et al., 1996). As a transcription factor, NHR-49 might directly regulate the expression of ODR-7 and/or ODR-10, so we tested the expression of these genes. Interestingly, although we observed no decrease in global (whole-worm) *odr-7* or *odr-10* mRNA expression, analysis of a transcription ODR-10::GFP reporter revealed a striking reduction of signal in *nhr-49* null mutants. These results suggest that either development, differentiation, or function of the AWA neurons may be compromised by *nhr-49* loss, indicating that this transcription factor, widely viewed as a regulator of physiology and metabolism, may exert more direct control over neuronal cells. The observed defect could arise either from cell-autonomous function or from indirect signals, which may include sphingolipids. Interestingly, sphingolipids play key roles in the nervous systems of many animals, including in aversive learning in *C. elegans* (Wu et al., 2025), and these roles include inter-tissue communication (Kim and Sieburth, 2018; Wang et al., 2023). Ultimately, further experiments will be required to dissect the mechanisms, cell autonomous or not and possible including cell signaling via lipids, through which NHR-49 promotes attractive chemotaxis.

It is noteworthy that ODR-7 itself encodes a divergent orphan nuclear hormone receptor (Colosimo et al., 2003). Little is known on how ODR-7 regulates the expression of its genes, but like many other NHRs (Taubert et al., 2011), it may form heterodimers with other NHRs. It is intriguing to speculate that ODR-7 may bind NHR-49 and thus achieve regulation of specific sets of mRNAs in AWA neurons to promote pertinent chemoattractant signaling. However, the broader role of NHR-49 in mechanosensation, which isn’t linked to AWA neurons and the fact that NHR-49 action from body wall muscle is sufficient to achieve diacetyl attraction argues against such a cell-type autonomous mechanism.

### NHR-49 promotes mechanosensation and short-term learning

Our data suggests that NHR-49 may play broader roles in neuronal sensing and signal transduction circuits than previously appreciated, leading us to expand our studies into the pathways governing mechanosensation and habituation. As a basic form of non-associative learning, habituation is a critical fundamental behavior observed in many higher organisms and is thought to be involved in the basis of more complex behaviors (Rose and Rankin, 2001). *C. elegans* display habituation after exposure to repetitive mechanosensory stimuli, and this behavior is controlled by a complex neural circuit. Typically, worms reverse after a mechanosensory stimulus to move away from a potential threat (Rose and Rankin, 2001). After repeated stimuli, the circuit regulating forward motion, driven by the PLM neuron in the tail, is activated, whereas the regulation of backward motion, driven by the AVM and ALM neurons in the midbody, is inhibited (Rose and Rankin, 2001). Thus, the population probability of reversal after repetitive mechanical stimuli decreases over time in wild-type worms, representing habituation. However, we found that *nhr-49* mutants retain stronger behavioral responses to mechanical stimuli over time, representing less pronounced habituation than wild-type worms.

NHR-49 is important in serotonergic neurons to regulate aversive behavior during infection (Kwon et al., 2024). The AWA neuron is not known to release any neurotransmitters and instead fires action potentials, while the neurons that regulate habituation vary in the neurotransmitters they use (Lee et al., 1999; Liu et al., 2018). Future experiments to dissect the roles of NHR-49 in each of these behaviors could provide clarity on whether NHR-49 acts via a common pathway to promote general neuron function or via independent pathways in each of these behaviors.

In sum, our studies reveal new neuronal behaviours that are under the control of NHR-49 independently of its known roles in lipid metabolism. Taken together with published studies, our data suggest that NHR-49 may integrate metabolic and behavioural responses across tissues to coordinate energy intake and metabolism. More detailed mapping of NHR-49 dependent gene expression in single cells and tissues may reveal these pathways in greater detail in the future.

## Acknowledgments

This work was supported by grants from the Canadian Institutes of Health Research (CIHR; PJT-153199, PJT-186144 to S.T., PJT-195874 to C.H.R) and the Natural Sciences and Engineering Research Council of Canada (NSERC; RGPIN-2018-05133 and RGPIN-2024-06537 to S.T.). G.E. was supported by CIHR CGS-D and UBC Medical Genetics Graduate Program scholarships, J.Y. by BCCHR and UBC scholarships, J.T.W.W. by NSERC Undergraduate Summer Research Award (USRA) and UBC Edwin S.H. Leong Centre for Healthy Aging Summer studentships, Y.W. by a BC Graduate Scholarship, and S.T. by a BCCHR Investigator Grant Award Program (IGAP) award. Some strains were provided by the CGC, which is funded by NIH Office of Research Infrastructure Programs (P40 OD010440).

## Author Contributions

Designed research: JTWW, GE, JY, LDK, MEZ, YL, CHR, ST; Performed research: JTWW, GE, JY, LDK, MEZ, YKOW, YL; Contributed unpublished reagents/ analytic tools: JY; Analyzed data: JTWW, GE, JY, LDK, MEZ, YKOW, YL; Wrote the paper: JTWW, GE, JY, LDK, MEZ, YKOW, YL, CHR, ST.

## References

Bargmann CI, Hartwieg E, Horvitz HR (1993) Odorant-selective genes and neurons mediate olfaction in C. elegans. Cell 74:515–527.

Brandel-Ankrapp KL, Arey RN (2023) Uncovering novel regulators of memory using C. elegans genetic and genomic analysis. Biochem Soc Trans 51:161–171.

Brenner S (1974) The genetics of Caenorhabditis elegans. Genetics 77:71–94.

Chiba CM, Rankin CH (1990) A developmental analysis of spontaneous and reflexive reversals in the nematode Caenorhabditis elegans. J Neurobiol 21:543–554.

Colbert HA, Bargmann CI (1995) Odorant-specific adaptation pathways generate olfactory plasticity in C. elegans. Neuron 14:803–812.

Colosimo ME, Tran S, Sengupta P (2003) The divergent orphan nuclear receptor ODR-7 regulates olfactory neuron gene expression via multiple mechanisms in Caenorhabditis elegans. Genetics 165:1779–1791.

Dasgupta M, Shashikanth M, Gupta A, Sandhu A, De A, Javed S, Singh V (2020) NHR-49 Transcription Factor Regulates Immunometabolic Response and Survival of Caenorhabditis elegans during Enterococcus faecalis Infection. Infect Immun 88.

Deline ML, Vrablik TL, Watts JL (2013) Dietary supplementation of polyunsaturated fatty acids in Caenorhabditis elegans. J Vis Exp 50879.

Doering KR, Cheng X, Milburn L, Ratnappan R, Ghazi A, Miller DL, Taubert S (2022) Nuclear Hormone Receptor NHR-49 acts in parallel with HIF-1 to promote hypoxia adaptation in Caenorhabditis elegans. Elife 11.

Doering KRS, Ermakova G, Taubert S (2023) Nuclear hormone receptor NHR-49 is an essential regulator of stress resilience and healthy aging in Caenorhabditis elegans. Front Physiol 14:1241591.

Ferkey DM, Sengupta P, L’Etoile ND (2021) Chemosensory signal transduction in Caenorhabditis elegans. Genetics 217:iyab004.

Gilst MRV, Hadjivassiliou H, Jolly A, Yamamoto KR (2005a) Nuclear Hormone Receptor NHR-49 Controls Fat Consumption and Fatty Acid Composition in C. elegans. PLoS biology 3:e53.

Gilst MRV, Hadjivassiliou H, Yamamoto KR (2005b) A Caenorhabditis elegans nutrient response system partially dependent on nuclear receptor NHR-49. Proc Natl Acad Sci USA 102:13496–13501.

Goodman MB, Sengupta P (2019) How Caenorhabditis elegans Senses Mechanical Stress, Temperature, and Other Physical Stimuli. Genetics 212:25–51.

Goudeau J, Bellemin S, Toselli-Mollereau E, Shamalnasab M, Chen Y, Aguilaniu H (2011) Fatty Acid Desaturation Links Germ Cell Loss to Longevity Through NHR-80/HNF4 in C. elegans. PLoS biology 9:e1000599.

Hammarlund M, Hobert O, Miller DM, Sestan N (2018) The CeNGEN Project: The Complete Gene Expression Map of an Entire Nervous System. Neuron 99:430–433.

Hilliard MA, Bargmann CI, Bazzicalupo P (2002) C. elegans Responds to Chemical Repellents by Integrating Sensory Inputs from the Head and the Tail. Curr Biol 12:730–734.

Kahn-Kirby AH, Dantzker JLM, Apicella AJ, Schafer WR, Browse J, Bargmann CI, Watts JL (2004) Specific polyunsaturated fatty acids drive TRPV-dependent sensory signaling in vivo. Cell 119:889–900.

Kepler LD, McDiarmid TA, Rankin CH (2022) Rapid assessment of the temporal function and phenotypic reversibility of neurodevelopmental disorder risk genes in C. elegans. Dis Model Mech 15:dmm049359.

Kim S, Sieburth D (2018) Sphingosine Kinase Regulates Neuropeptide Secretion During the Oxidative Stress-Response Through Intertissue Signaling. J Neurosci 38:8160–8176.

Kurashina M, Mizumoto K (2023) Targeting endogenous proteins for spatial and temporal knockdown using auxin-inducible degron in Caenorhabditis elegans. STAR Protoc 4:102028.

Kwon S, Park K-S, Yoon K (2024) Regulator of Lipid Metabolism NHR-49 Mediates Pathogen Avoidance through Precise Control of Neuronal Activity. Cells 13:978.

Kwon S, Park K-S, Yoon K (2023) Dissecting the Neuronal Contributions of the Lipid Regulator NHR-49 Function in Lifespan and Behavior in C. elegans. Life 13:2346.

Lee RYN, Sawin ER, Chalfie M, Horvitz HR, Avery L (1999) EAT-4, a Homolog of a Mammalian Sodium-Dependent Inorganic Phosphate Cotransporter, Is Necessary for Glutamatergic Neurotransmission in Caenorhabditis elegans. J Neurosci 19:159–167.

Liu Q, Kidd PB, Dobosiewicz M, Bargmann CI (2018) C. elegans AWA Olfactory Neurons Fire Calcium-Mediated All-or-None Action Potentials. Cell 175:57–70.e17.

Margie O, Palmer C, Chin-Sang I (2013) C. elegans chemotaxis assay. Journal of visualized experiments: JoVE e50069.

Naim N, Amrit FRG, Ratnappan R, DelBuono N, Loose JA, Ghazi A (2021) Cell nonautonomous roles of NHR 49 in promoting longevity and innate immunity. Aging Cell e13413.

Nuttley WM, Harbinder S, Kooy D van der (2001) Regulation of Distinct Attractive and Aversive Mechanisms Mediating Benzaldehyde Chemotaxis in Caenorhabditis elegans. Learn Mem 8:170–181.

O’Halloran DM, Altshuler-Keylin S, Lee JI, L’Etoile ND (2009) Regulators of AWC-Mediated Olfactory Plasticity in Caenorhabditis elegans. PLoS Genet 5:e1000761.

Pathare PP, Lin A, Bornfeldt KE, Taubert S, Van Gilst MR (2012) Coordinate regulation of lipid metabolism by novel nuclear receptor partnerships. PLoS genetics 8:e1002645.

Pender CL, Horvitz HR (2018) Hypoxia-inducible factor cell non-autonomously regulates C. elegans stress responses and behavior via a nuclear receptor. eLife 7:e36828.

Rankin CH, Beck CDO, Chiba CM (1990) Caenorhabditis elegans: A new model system for the study of learning and memory. Behav Brain Res 37:89–92.

Ratnappan R, Amrit FRG, Chen S-W, Gill H, Holden K, Ward J, Yamamoto KR, Olsen CP, Ghazi A (2014) Germline Signals Deploy NHR-49 to Modulate Fatty-Acid β-Oxidation and Desaturation in Somatic Tissues of C. elegans. Plos Genet 10:e1004829.

Rose JK, Rankin CH (2001) Analyses of habituation in Caenorhabditis elegans. Learn Mem 8:63–69.

Sabatella M, Thijssen KL, Davó-Martínez C, Vermeulen W, Lans H (2021) Tissue-Specific DNA Repair Activity of ERCC-1/XPF-1. Cell Rep 34:108608.

Sengupta P, Chou JH, Bargmann CI (1996) odr-10 Encodes a Seven Transmembrane Domain Olfactory Receptor Required for Responses to the Odorant Diacetyl. Cell 84:899–909.

Sengupta P, Colbert HA, Bargmann CI (1994) The C. elegans gene odr-7 encodes an olfactory-specific member of the nuclear receptor superfamily. Cell 79:971–980.

Swierczek NA, Giles AC, Rankin CH, Kerr RA (2011) High-Throughput Behavioral Analysis in C. elegans. Nat Methods 8:592–598.

Tatge L, Douglas PM (2025) Isoform differences drive functional diversity of NHR-49. MicroPubl Biol 2025.

Taubert S, Ward JD, Yamamoto KR (2011) Nuclear hormone receptors in nematodes: evolution and function. Molecular and cellular endocrinology 334:49–55.

Tsai S-H, Wu Y-C, Palomino DF, Schroeder FC, Pan C-L (2024) Peripheral peroxisomal β-oxidation engages neuronal serotonin signaling to drive stress-induced aversive memory in C. elegans. Cell Rep 43:113996–113996.

Van der Auwera P, Frooninckx L, Buscemi K, Vance RT, Watteyne J, Mirabeau O, Temmerman L, De Haes W, Fancsalszky L, Gottschalk A, Raizen DM, Nelson MD, Schoofs L, Beets I (2020) RPamide neuropeptides NLP-22 and NLP-2 act through GnRH-like receptors to promote sleep and wakefulness in C. elegans. Sci Rep 10:9929.

Wang W, Sherry T, Cheng X, Fan Q, Cornell R, Liu J, Xiao Z, Pocock R (2023) An intestinal sphingolipid confers intergenerational neuroprotection. Nat Cell Biol 25:1196–1207.

Wani KA, Goswamy D, Taubert S, Ratnappan R, Ghazi A, Irazoqui JE (2021) NHR-49/PPAR-α and HLH-30/TFEB cooperate for C. elegans host defense via a flavin-containing monooxygenase. eLife 10.

Watts JL, Ristow M (2017) Lipid and Carbohydrate Metabolism in Caenorhabditis elegans. Genetics 207:413–446.

Wu Y-C, Beets I, Fox BW, Palomino DF, Chen L, Liao C-P, Vandewyer E, Lin L-Y, He C-W, Chen L-T, Lin C-T, Schroeder FC, Pan C-L (2025) Intercellular sphingolipid signaling mediates aversive learning in C. elegans. Curr Biol 35:2323–2336.e9.

Yan J, Bhanshali F, Shuzenji C, Mendenhall TT, Taylor SKB, Ermakova G, Cheng X, Bai P, Diwan G, Seraj D, Meyer JN, Sorensen PH, Hartman JH, Taubert S (2025) Eukaryotic Elongation Factor 2 Kinase EFK-1/eEF2K promotes starvation resistance by preventing oxidative damage in C. elegans. Nat Commun 16:1752.

Yokosawa R, Noma K (2025) A Nuclear Hormone Receptor nhr 76 Induces Age Dependent Chemotaxis Decline in C. elegans. Aging Cell 24:e70277.

Zhang L, Ward JD, Cheng Z, Dernburg AF (2015) The auxin-inducible degradation (AID) system enables versatile conditional protein depletion in C. elegans. Development 142:4374–4384.

Zhang Y, Iino Y, Schafer WR (2024) Behavioral plasticity. GENETICS 228:iyae105.

